# *RadiationGeneSigDB*: A database of oxic and hypoxic radiation response gene signatures and their utility in identification of hypoxia-regulated MicroRNA

**DOI:** 10.1101/455345

**Authors:** Venkata SK. Manem, Andrew Dhawan

**Author notes:** **Corresponding Author:** Venkata SK Manem, Ph.D. Research Professional Institut Universitaire de Cardiologie et de Pneumologie de Québec, Université Laval, QC, Canada.

## Abstract

**Summary:** Radiation therapy is among the most effective and widely used modalities of cancer therapy in current clinical practice. With the advent of new high throughput genomic technologies and the continuous inflow of transcriptomic data, there has been a paradigm shift in the landscape of radiation oncology. In this era of personalized radiation medicine, genomic datasets hold great promise to investigate novel biomarkers predictive of radiation response. In this regard, the number of available gene expression based signatures built under oxic and hypoxic conditions is getting larger. This poses two main questions in the field, namely, *i)* how reliable are these signatures when applied across a compendium of datasets in different model systems; and *ii)* is there redundancy of gene signatures. To address these fundamental radiobiologic questions, we curated a database of gene expression signatures predictive of radiation response under oxic and hypoxic conditions. *RadiationGeneSigDB* has a collection of 11 oxic and 24 hypoxic signatures with the standardized gene list as a gene symbol, Entrez gene ID, and its function. We present the utility of this database through three case studies: *i)* comparing breast cancer oxic signatures in cell line data *vs.* patient data; *ii)* comparing the similarity of head and neck cancer hypoxia signatures in clinical tumor data; and *iii)* gaining an understanding of hypoxia-associated miRNA. This valuable, curated repertoire of published gene expression signatures provides a motivating example for how to search for similarities in radiation response for tumors arising from different tissues across model systems under oxic and hypoxic conditions, and how a well-curated set of gene signatures can be used to generate novel hypotheses about the functions of non-coding RNA.

**Availability and implementation:** *RadiationGeneSigDB* is implemented in R. The source code of this package and signatures can be downloaded from the GitHub: https://github.com/vmsatya/RadiationGeneSigDB

## 1 Introduction

The two pillars driving the field of personalized radiation oncology are: *i)* treatment delivery and dose conformity arising from technological improvements, which include particle therapies and advanced image guidance techniques; *ii)* novel biomarker-guided tools, integrating concomitant chemotherapy [1]. To tailor radiation therapy, it is crucial to build predictive assays that are more confidently able to stratify patients, and concomitantly have associated impactful radiotherapeutic regimens. This could augment the existing radiobiological treatment strategies to more biologically-driven personalized radiation treatment to individual patients. The evolution of high throughput technologies and the continuous inflow of transcriptomic data have created new avenues to understand complex biological events induced by radiation, through data driven analysis, at a level beyond the gross clinical variables of an individual, and instead, at the individual tumor level. The ever-expanding arsenal of transcriptomic data holds great promise to investigate novel biomarkers that are predictive of radiation response. In the literature, several studies have attempted to associate radiosensitivity with molecular/genomic features [2], and this number is getting larger. But, there have been no systematic efforts to build a database of *radiation response* gene expression signatures validated or designed for clinical use.

In the literature, many groups conducted comprehensive gene expression profiling and built signatures for radiation response under oxic and hypoxic conditions. Two methods have been used to identify these gene signatures, namely, data-driven (bottom-up) and hypothesis-based (top-down) approaches. The performances of these transcriptomic signatures have been evaluated on various datasets with limited to no independent validation. Moreover, there is only minimal overlap between these gene signatures. This may be attributed to the different platforms, such as microarray or RNA-sequencing, training sets, and statistical tools used to build these signatures. Furthermore, in order to develop biomarkers reproducible and appropriate for clinical translation, a database should be built to house these predictive models. At present, there is no radiation response signature database that could potentially address these fundamental questions. A wealth of molecular data [3,4] and gene expression signatures [5] are publicly available through diverse online resources. In this study, we manually curated a number of radiation response gene signatures from the literature, known as the *RadiationGeneSigDB*, and implemented this as an *R* package. This database will facilitate users, *i)* to compare radiation response signatures across pan-cancer datasets; *ii)* to investigate the prognostic value of these signatures on a compendium of clinical datasets using meta-analysis; and, *iii)* to investigate the tissue specificity of radiation response using these signatures across different transcriptomic platforms. Ultimately, the goal of this work is to improve the development, and spur a greater understanding of how transcriptomic signatures can be used to augment precision radiation oncology, for improved patient outcomes, and more effectively designed clinical trials.

## 2 Methods

### 2.1 Curation of RadiationGeneSigDB

In most online resources, gene signatures are often included in tables or figures embedded in publications. These signatures often use non-standard gene identifiers, making comparison to other gene signatures, or even to the original data a significant challenge. To be of maximal value, these gene signatures should be available through an easily accessible database resource that provides gene sets in a more standard format. Radiation response gene expression based signatures are identified in Pubmed. In our first release, we collected 11, 24 oxic and hypoxic signatures respectively, and these are described in Supplementary Tables 1 and 2. We manually annotated them with gene symbol, Entrez Gene ID, and gene function, which can be downloaded from the GitHub as an excel file, or as an *R* object. The molecular subtyping for breast cancer, computation of signature score and the quality control of signatures used in this study is discussed in the Supplementary methods.

**Table 1:** miRNA significantly positively and negatively associated with hypoxia gene signatures (top 10 displayed). miRNA are identified as significantly associated by way of linear modelling schema, as outlined in the methods section, and by considering the association between miRNA expression and hypoxia gene signature score across each of the 24 gene signatures curated into *RadiationGeneSigDB*.

|  |  |
| --- | --- |
| miRNA positively associated with hypoxia | miRNA negatively associated with hypoxia |

**Table 1:** miRNA significantly positively and negatively associated with hypoxia gene signatures (top 10 displayed).
| miRNA | Rank Product p value | miRNA | Rank product p value |
| --- | --- | --- | --- |
| hsa-miR-210-3p | 3.15E-42 | hsa-miR-374a-5p | 2.83E-31 |
| hsa-miR-15b-5p | 1.98E-27 | hsa-miR-29c-5p | 3.04E-21 |
| hsa-miR-130b-5p | 1.03E-21 | hsa-miR-1307-3p | 1.91E-18 |
| hsa-miR-93-5p | 2.38E-18 | hsa-miR-30e-3p | 4.23E-15 |
| hsa-miR-21-3p | 5.99E-16 | hsa-miR-30e-5p | 1.46E-14 |
| hsa-miR-106b-5p | 1.94E-10 | hsa-let-7i-3p | 1.65E-12 |
| hsa-miR-21-5p | 2.96E-10 | hsa-miR-26a-5p | 1.12E-11 |
| hsa-miR-223-3p | 3.15E-10 | hsa-let-7a-3p | 1.27E-11 |
| hsa-miR-15b-3p | 9.14E-10 | hsa-miR-362-5p | 8.40E-11 |

### 2.2 Datasets

Cancer cell lines were profiled at the genomic level and the processed data are available for download from a public database, The Cancer Cell Line Encyclopedia (CCLE) [6]. For this study, we used all breast cancer lines (26 in total) from the CCLE dataset. The METABRIC dataset was used for patient data [3] and selected only those patients treated with radiation therapy. Out of 1992 patients, 232 patients were reportedly treated with radiation. We retrieved head and neck squamous cell carcinoma (HNSCC) primary tumor transcriptomic data from the Tumor Cancer Genome Atlas (TCGA) [4], and selected only the patients treated with radiation (99 in total).

### 2.3 Predictive Model to Obtain Hypoxia-Associated miRNA

Genomic data involving the mRNA and miRNA expression of a cohort of 7,738 patient samples was downloaded from the Cancer Genome Atlas (TCGA) project, accessed through the Broad Institute Firebrowse portal at http://www.firebrowse.org [4]. Data used were RSEM-normalised gene expression and mature miRNA normalised expression. In the same method as Dhawan et al. [7], (In press, Nature Communications), we considered all cancer types which were epithelial or glandular with respect to histology, and with at least 200 unique patient samples with paired mRNA and miRNA-sequencing data (Supplementary Table 3).

In the same method of Dhawan et al., 2018, mRNA gene signature scores, scored as the median value of gene expression for the genes of a given signature, were taken. These values were used as the response variables in two series of predictive models. In the first, every miRNA with non-zero expression across at least 80% of all tumour samples was considered for its correlation with the gene signature score, across all samples of a given cancer type. miRNAs showing at least moderate univariate predictive ability for the signature summary score, were considered going forward. Subsequently, each of these miRNA passing the first filter were taken as predictors for a linear model, fitting the gene signature scores. Fitting was done by the same approach as taken by Dhawan et al. 2018, Nature Communications. That is, multivariable linear regression with L1/L2 penalty, optimized by 10-fold cross-validation was used to identify the miRNAs which showed the greatest predictive ability for each hypoxia gene signature score across each of the cancer types considered. miRNA which showed the strongest predictive ability, in positive or negative association with the gene signature score were obtained for each signature, by aggregating the linear model coefficients across cancer types, and considering the rank product statistic. Those miRNA that consistently had greater positive coefficients in each linear model across cancer types, than chance alone could predict, were taken as significantly positively associated to a given gene signature. These significantly-associated miRNA were then ranked in order of p-value obtained by rank product statistic, for each gene signature considered.

Subsequently, these miRNA were aggregated across all gene signatures, to identify that recurrently positively or negatively associated with each gene signature, using the rank product taken over all gene signatures.

## 3 Results

We present here three case studies utilizing the radiation response gene signature database, highlighting its utility, and the associated *R* code is provided in the Github.

### 3.1 Comparison of *oxic* breast gene signatures in cell lines and patients

The objective of this case study is to conduct an unbiased comparison of two different prognostic breast cancer signatures (namely, Piening et al. [8], Speers et al. [9]) predictive of radiation response under oxic conditions. We compared the signature scores in cell line data as well as in patients, and assessed the similarities and differences in their behavior across these datasets. Importantly, the overlap of genes between these two signatures is minimal, attributable to the different platforms, training sets, and statistical methods used in their generation. We computed signature scores across the 26 breast cancer cell lines from the the CCLE RNA-seq database, as well as for all patient samples from the METABRIC dataset. Before proceeding further with this analysis, we first assayed the quality of gene signature application on each of these datasets using *sigQC*. Interestingly, this precursor analysis, with summary radar plots (Figure 1), showed that there are significant variations in quality between the signatures on different datasets. That is, the signatures appeared to have strongest quality when applied on CCLE data or TNBC data, but not the ER-low subtype.

**Figure 1:**
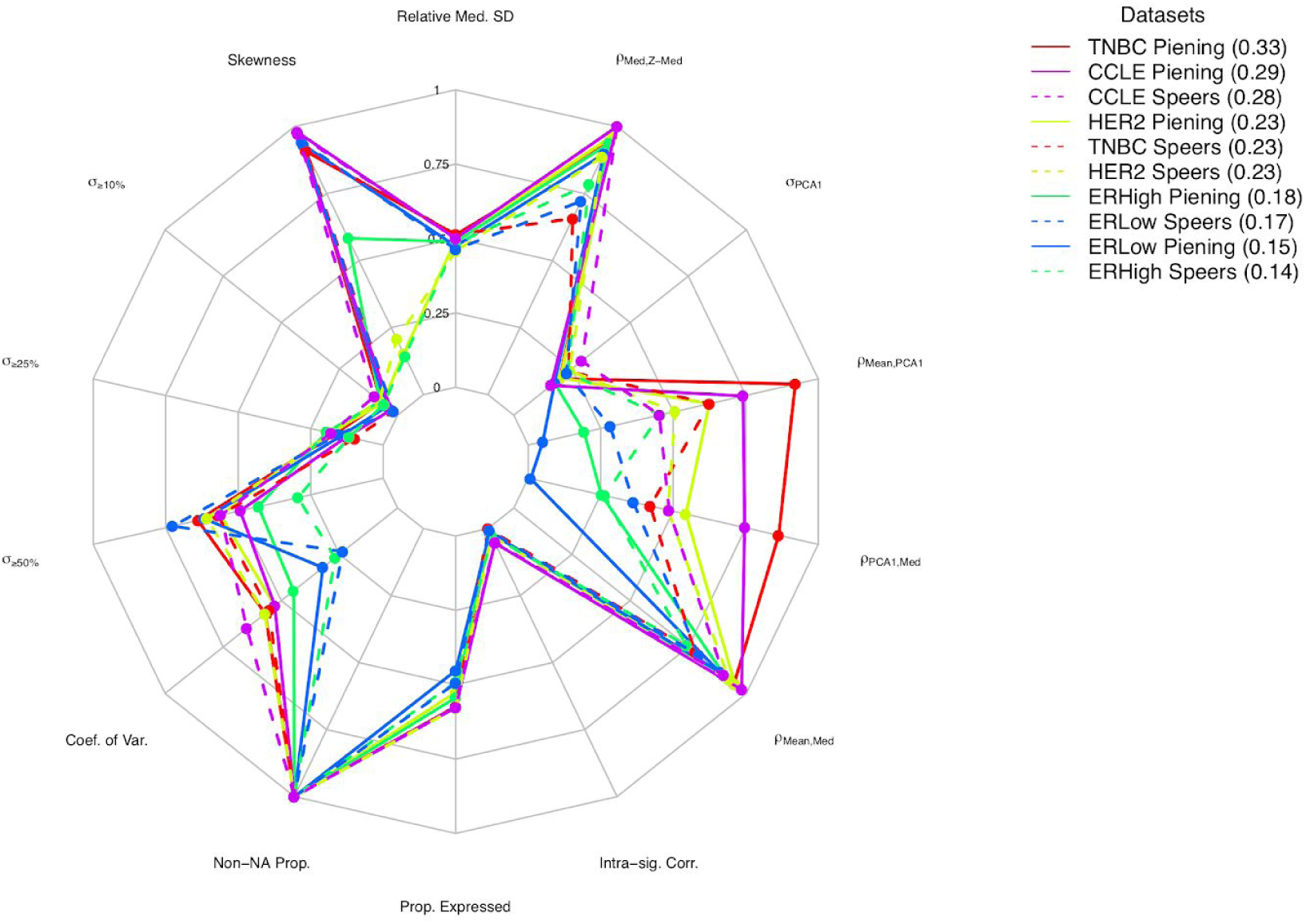
Radar plots produced for *sigQC* summary metric evaluation of Piening and Speers signatures on CCLE and METABRIC datasets. Each ray of the radar plot evaluates one of the summary metrics checked by *sigQC* for gene signature quality prior to application on a dataset.

Briefly, this plot reveals that the Piening signature on the TNBC dataset (solid red line) is among the highest-quality application, along with the Piening signature on the CCLE dataset (solid purple line). Among the lowest performing signature-dataset combinations are the Speers and Piening signatures on the ER-High and ER-Low datasets, as poorer performance is observed on nearly every metric considered. Signature-dataset combinations are ranked in the legend by quality, with the numeric value in brackets representing the area contained within the radar chart lines for each signature-dataset combination.

Next, the signature scores for both the Piening and Speers signatures were then compared directly with the Spearman correlation coefficient (Figure 2). We observed a moderate Spearman correlation (ρ ~0.69) between the signature scores in the cell line data. However, we observed significant differences when comparing signature scores in patient data. We observed that for clinical data, when stratified by molecular subtype, the Spearman correlation has moderate value for triple negative breast cancer, HER2 and ER-low cancers. For the ER-high subtype, the Spearman correlation between the signatures is weak (ρ ~0.25). The moderate correlation between the two breast cancer signatures can be attributed to: *i)* different statistical tools implemented on different platforms, with a small overlap in genes; *ii)* different experimental assays and protocols used to generate dose-response data; *iii)* inconsistent experimental methodologies to generate radiation data across labs; *iv)* signature development underpowered due to lack of sample size along with enough replicates. This could also have implications with regard to the evaluation of radiation response in an *in-vitro* setting and translating it to the clinical practice. Hence, there is a dire need to build robust radiation signatures that are independent of platforms and technology.

**Figure 2:**
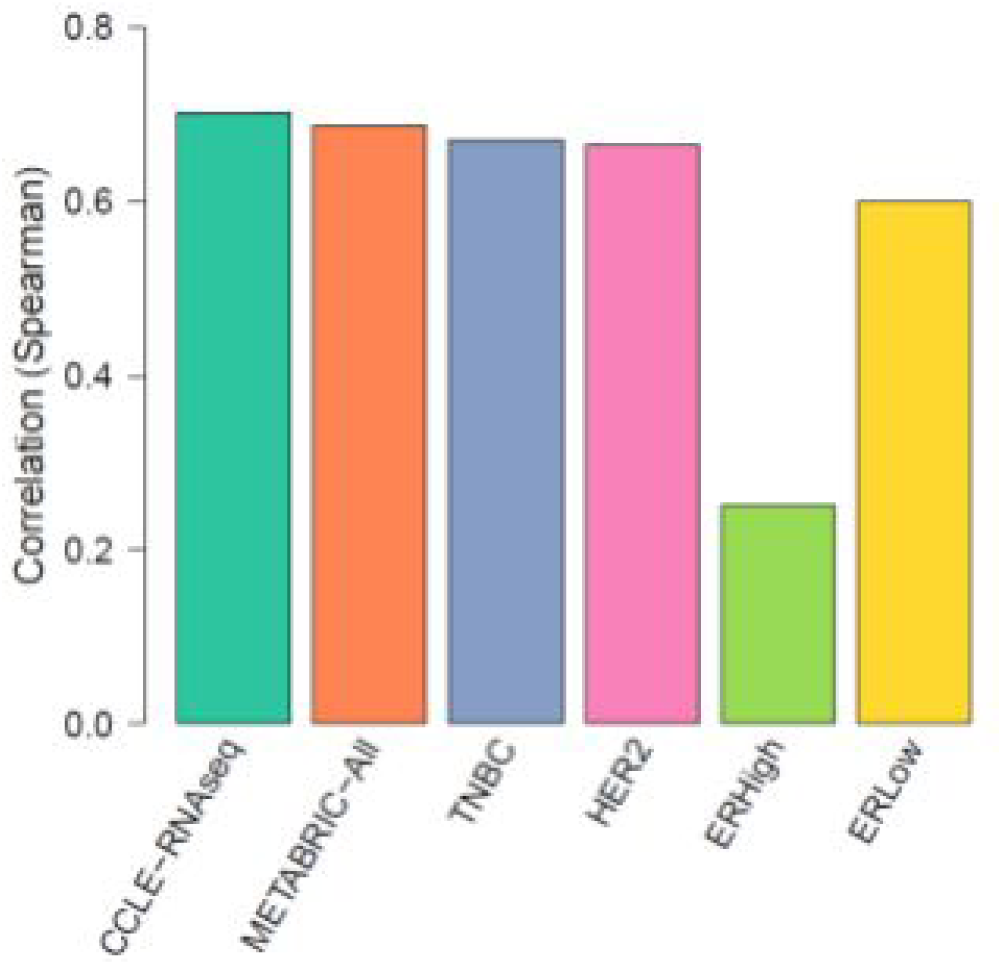
Cell line vs. Patient model systems. Comparison of breast cancer radiation response signature scores in CCLE breast cancer cell lines and in METABRIC patient cohort.

### 3.2 Comparison of HNSCC *hypoxia* gene signatures in patients

In this case study, we aimed to compare three HNSCC hypoxia gene expression signatures in the TCGA cohort. The first hypoxia signature is identified by Toustrup et al. [10], which was validated in a HNSCC cohort. The second hypoxia signature is developed by Eustace et al. [11] on laryngeal tumors, which was validated in HNSCC cohort. Lastly, the third signature was built by Lendahl et al. [12], which was derived to determine a pan-cancer signature for radiation response in hypoxia. Firstly, comparing the composition of each of these signatures, we noted that there is just a single common gene between the three signatures. Next, we performed a quality analysis, to ensure legitimacy when applying these signatures on the TCGA dataset with *sigQC* and noted that all three signatures show good quality on this dataset and are consistent between each other (Figure 3).

**Figure 3:**
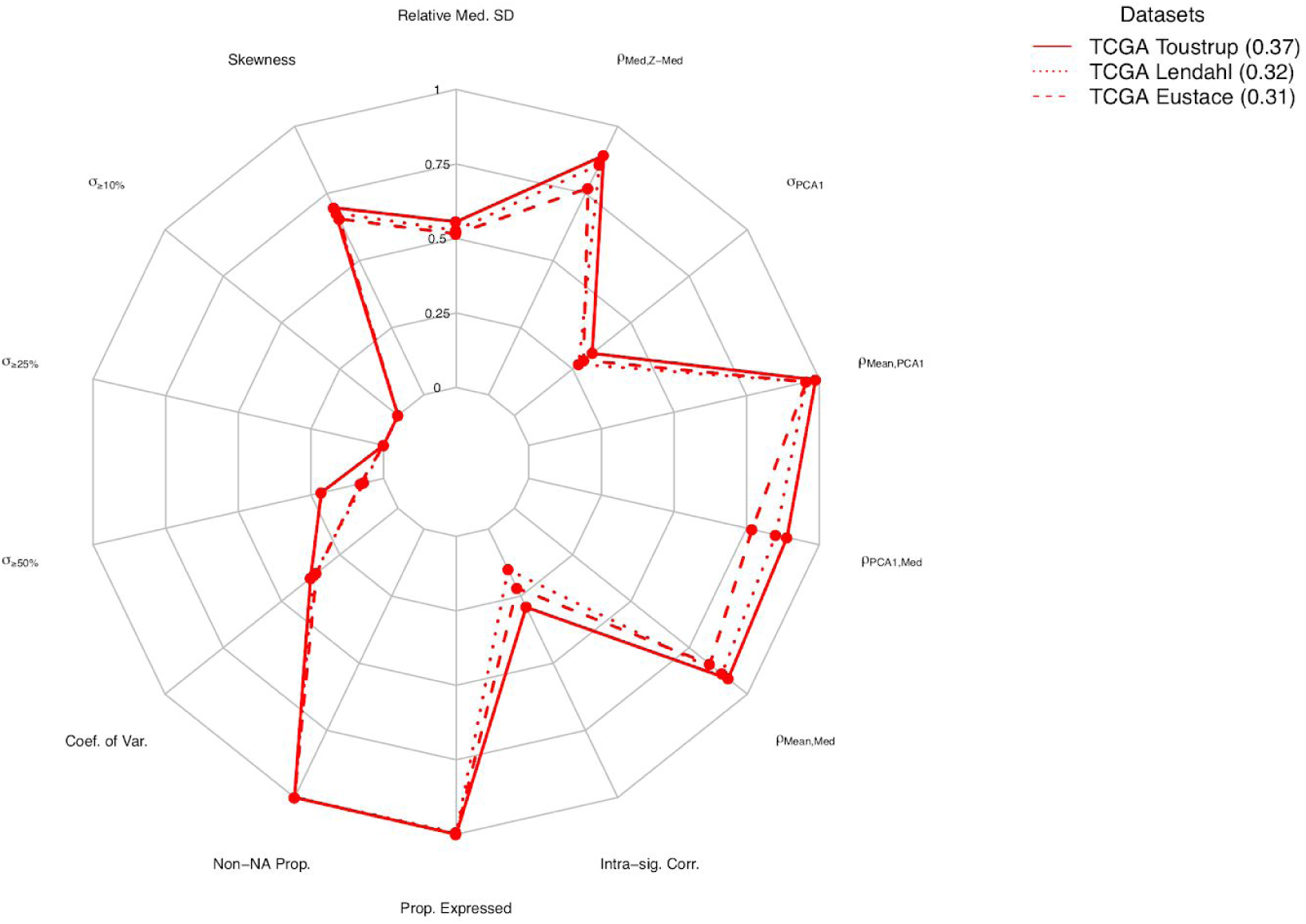
Radar plots produced for *sigQC* summary metric evaluation of Toustrup, Lendahl, and Eustace signatures on TCGA HNSCC dataset. Each ray of the radar plot evaluates one of the summary metrics checked by *sigQC* for gene signature quality prior to application on a dataset.

Briefly, this plot reveals that each of the three signatures has approximately similar quality when applied on this dataset, with the Eustace signature showing a slight reduction in quality, owing primarily to differences in signature gene score correlation, likely owing to model-specificity in its use. Signature-dataset combinations are ranked in the legend by quality, with the numeric value in brackets representing the area contained within the radar chart lines for each signature-dataset combination. We found that the Toustrup and Lendahl, and the Toustrup and Eustace hypoxia signatures were strongly correlated (ρ ~0.8), whereas the Lendahl and Eustace signatures were weakly correlated (ρ ~0.46) (Figure 4-Left panel), which is consistent with [13]. Having identified differences in their genetic composition, but relative similarity in the behavior, we next asked whether similar biological processes could be defining these signatures. To achieve this, we performed pathway analysis using the GO terms from the MSigDB [5]. For an FDR < 10%, 13, 37, and 51 transcriptional pathways were found to be enriched using Toustrup, Eustace, Lendahl signatures respectively. We found only 3 pathways that were commonly enriched between all the 3 signatures (Figure 2-Right panel), namely, GO:Oxidation Reduction Process”, “GO:Glucose Metabolic Process,” GO:Monosaccharide Metabolic Process. We found 2 pathways to be enriched related to oxygen levels using Toustrup and Lendahl signatures, but not enriched using the Eustace signature (Figure 4-Right panel).

**Figure 4:**
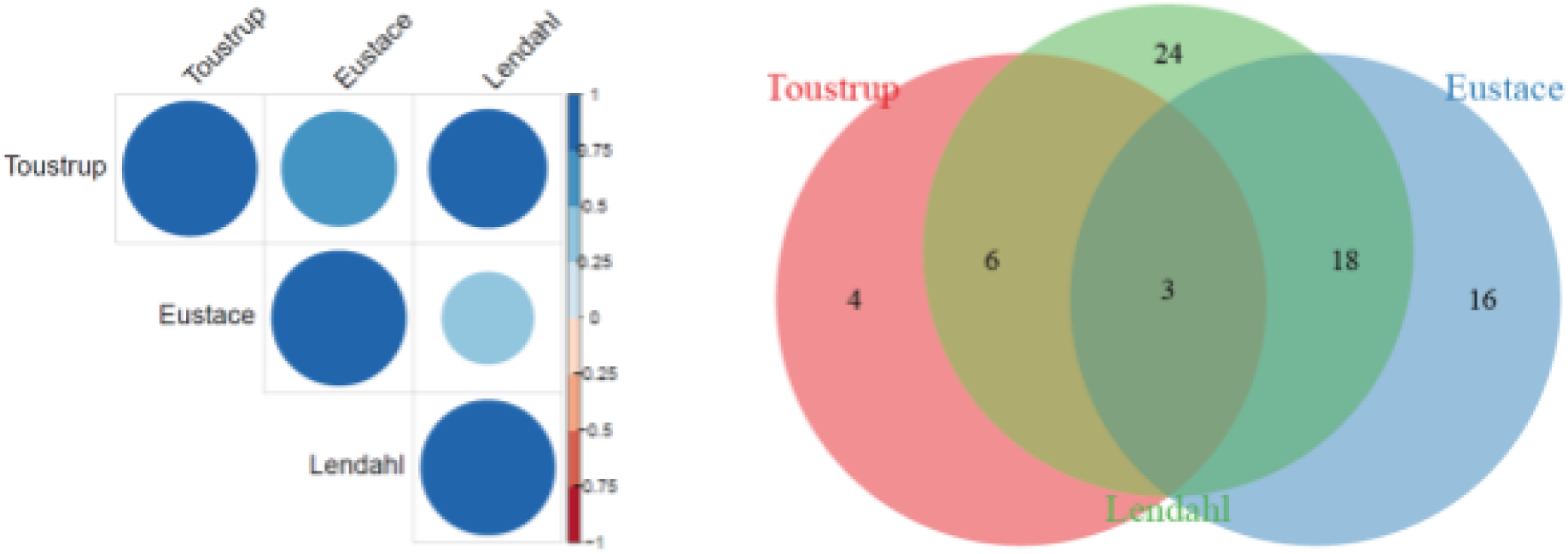
Comparison of hypoxia gene signatures in TCGA HNSCC. Left panel: Correlation of signature scores. **Right panel:** Venn diagram illustrating the transcriptional pathways enriched using the three head and neck hypoxia signatures (FDR<10%).

### 3.3 Hypoxia gene signatures used to identify hypoxia-associated miRNA

mRNA gene signatures, as generated over the past decade have enabled a greater understanding of cellular processes and phenotypic responses to environmental changes, such as hypoxia. Using *RadiationGeneSigDB*, the curated hypoxia gene signatures at the mRNA level were used to infer the function of miRNA using a robust pan-cancer statistical approach. Briefly, as described in the methods section, a graduated linear modelling approach with a pan-cancer dataset was used to identify those miRNA with significant and recurring statistical association to the overall behaviour of the hypoxia gene signatures. Using a strong, broadly-representative, and curated database of hypoxia gene signatures is essential to this approach, as the strength of the miRNA associations determined through the linear model.

As such, using the curated database of 24 hypoxia gene signatures and data from patient samples from 15 epithelial tumour types, we obtain a set of miRNA highly associated both positively and negatively to these hypoxia gene signatures. As a result of both the high quality of the signatures considered, as well as the robust statistical approach considered, the miRNA obtained as signature-associated include many of those that have been already independently validated, as summarised in Table 1, and as summarised in Figure 5. Importantly, the strength of this approach is its versatility - with any such well-curated list of gene signatures, or indeed a different set of genomic data, novel associations of miRNA to phenotype can be discovered, underlying the utility of this resource as a tool for biological discovery.

**Figure 5:**
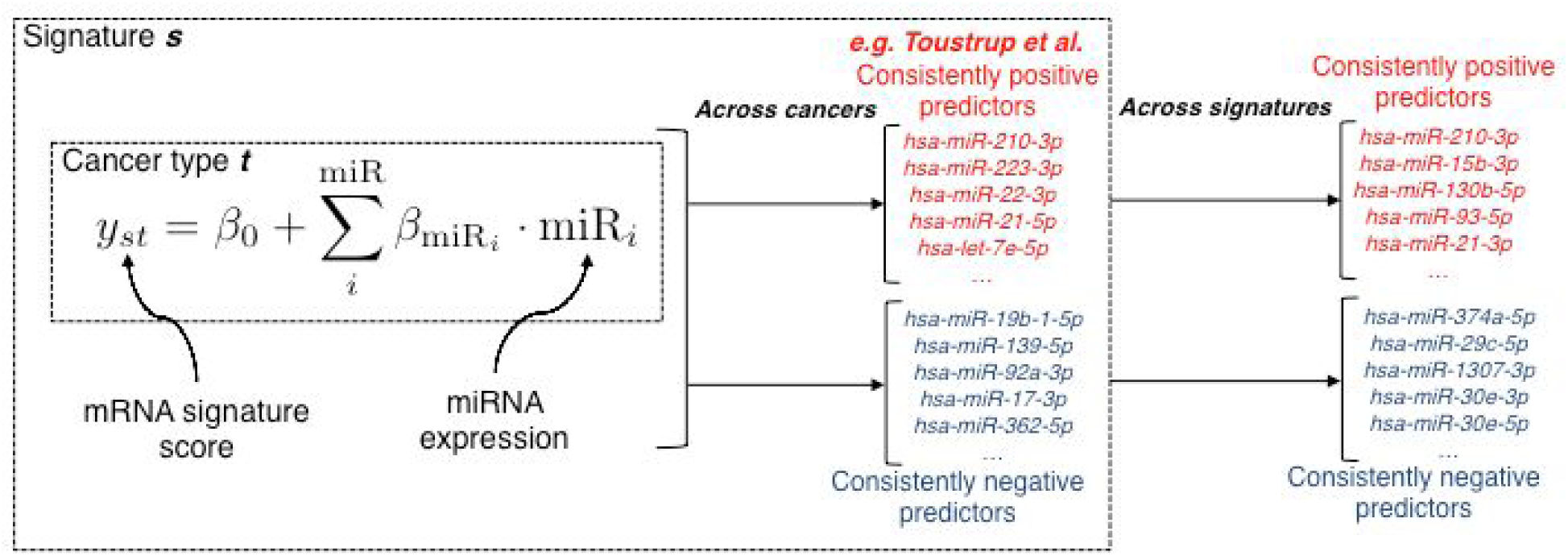
Overview of approach used to identify hypoxia-associated miRNA; figure adapted from Dhawan et al., Nature Communications. Figure depicts on left side an overview of the linear model used in fitting, wherein each gene signature and cancer type are considered, and the miRNA significantly associated with a given signature across cancer types are stored. Subsequently, those miRNA which associate the strongest with all gene signatures are next considered before obtaining the final list of hypoxia-associated miRNA, as presented in Table 1.

## 4 Conclusions

In this era of personalized medicine, an area of current excitement in the field of radiation oncology is the study of changes in the transcriptome induced by radiation therapy, termed *radiogenomics*. With the evolution of sequencing technologies, radiogenomics has emerged as a new field to identify the biomarkers that are predictive of radiation response. Due to the continuous growth of transcriptomic studies across multiple cancers, the number of biomarkers predictive of radiation response is ever-increasing. The lack of genomic indicators of radiation response has impeded the administration of radiation to individual patients. Moreover, if a predictive biomarker were to be integrated into the clinical practice, it would require a reproducible assay that is agnostic to platforms and technologies. This study is an effort to build a database of radiation response gene signatures under *oxic* and *hypoxic* conditions. This repertoire of gene expression signatures are publicly available as an open source package in *R*. This will facilitate comparison of radiation response signatures across cancer types, and also enable us to investigate the prognostic value of these signatures using meta-analysis approaches. We hope and envision that this package will help users to compare their own signature to those in the *RadiationGeneSigDB* database, and help build better biomarkers by using multiple datasets in the discovery, or pre-clinical phase. Furthermore, *RadiationGeneSigDB* coupled with *RadioGx, a* novel computational platform of radiogenomics datasets [14], will enable us to build reliable and clinically-verifiable genomic predictors of radiation response.

## Availability of data and material

*RadiationGeneSigDB* is implemented in R. The source code of this package and signatures can be downloaded from the GitHub: https://github.com/vmsatya/RadiationGeneSigDB

## Competing interests

The authors declare that they have no competing interests.

## Funding

No funding is reported for this study

## Authors’ contributions

VM conceived the study, curated gene signatures from the literature. VM and AD performed the analysis, and interpreted the results. VM and AD wrote the manuscript. Both authors read and approved the final manuscript.

## Acknowledgements

The authors thank the scientific community for sharing their valuable data.

## Supplementary Material

### Supplementary Methods

We used the SCMOD2 model [1] to assign each tumor sample into the four established molecular subtypes of breast cancer: Basal-like (TNBC), Her2-enriched, Luminal A (ER-high) and Luminal B (ER-low). We used the SCMOD2 implementation available in the *genefu* R package. We computed a signed average (the sign being determined by the sign of the gene coefficient or direction) using the sig.score function in *genefu*. We computed the correlation between the the published gene signatures using the Spearman correlation coefficient. To assay the quality of gene signatures, we used our recently developed method *siqQC [2]*.

### Supplementary Tables

**Supplementary Table 1.** presents the original study, number of genes in the signature and the histology for the radiation response gene expression signatures under **oxic conditions** (fully oxygenated).

| Signature | Number of genes | Histology |
| --- | --- | --- |
| Piening et al. [3] | 281 | Breast Cancer |
| Nuyten et al. [4] | 335 | Breast Cancer |
| Torres-Roca et al. [5] | 10 | Pan-cancer |
| Speers et al. [6] | 51 | Breast Cancer |
| Pitroda et al. [7] | 4 | Carcinoma |
| Kim et al. [8] | 31 | Pan-cancer |
| De-Jong et al. [9] | 1177 | Head and Neck Squamous Cell Carcinoma |
| Amundson et al. [10] | 21 | Pan-cancer |
| Abazeed et al. [11] | 2248 | Squamous Cell Lung Cancer |
| Weichselbaum et al. [12] | 49 | Squamous Cell Carcinoma |
| Tang et al. [13] | 26 | Soft Tissue Sarcoma |

**Supplementary Table 2.** presents the original study, number of genes in the signature and the histology for the radiation response gene expression signatures under **hypoxic conditions** (absence of oxygen).

| **Signature** | **Number of genes** | **Histology** |
| --- | --- | --- |
| Toustrup et al. [14] | 15 | Head and Neck Cancer |
| Eustace et al. [15] | 26 | Head and Neck Cancer |
| Lendahl et al. [16] | 30 | Head and Neck Cancer |
| Winter et al. [17] | 99 | Pan-Cancer |
| Buffa et al. [18] | 51 | Pan-Cancer |
| Hu et al. [18,19] | 13 | Breast Cancer |
| Sorensen et al. [20] | 28 | Squamous Cell Carcinoma |
| Yang et al. [21] | 28 | Prostate Cancer |
| Fjeldbo CS et al. [22] | 6 | Cervical Cancer |
| Lang et al. [23] | 24 | Bladder Cancer |
| Ragnum et al. [24] | 32 | Prostate Cancer |
| Seigneuric et al. [25]<br>(Early 0% hypoxia) | 72 | Mammary Epithelial |
| Seigneuric et al. [25]<br>(Late 0% hypoxia) | 71 | Mammary Epithelial |
| Seigneuric et al. [25]<br>(Early 2% hypoxia) | 34 | Mammary Epithelial |
| Seigneuric et al. [25]<br>(Late 2% hypoxia) | 32 | Mammary Epithelial |
| Starmans et al. [26]<br>(Cluster 1) | 69 | Breast Cancer |
| Starmans et al. [26]<br>(Cluster 2) | 246 | Breast Cancer |
| Starmans et al. [26] | 157 | Breast Cancer |
| (Cluster 3) |  |  |
| Starmans et al. [26]<br>(Cluster 4) | 95 | Breast Cancer |
| Starmans et al. [26]<br>(Cluster 5) | 162 | Breast Cancer |
| Starmans et al. [26]<br>(Cluster 6) | 14 | Breast Cancer |
| Starmans et al. [26]<br>(Cluster 7) | 28 | Breast Cancer |
| Ghazoui et al. [27] | 70 | ER+ Breast Cancer |
| Van Malenstein et al. [28] | 7 | Hepatocellular Carcinoma |

**Supplementary Table 3.**
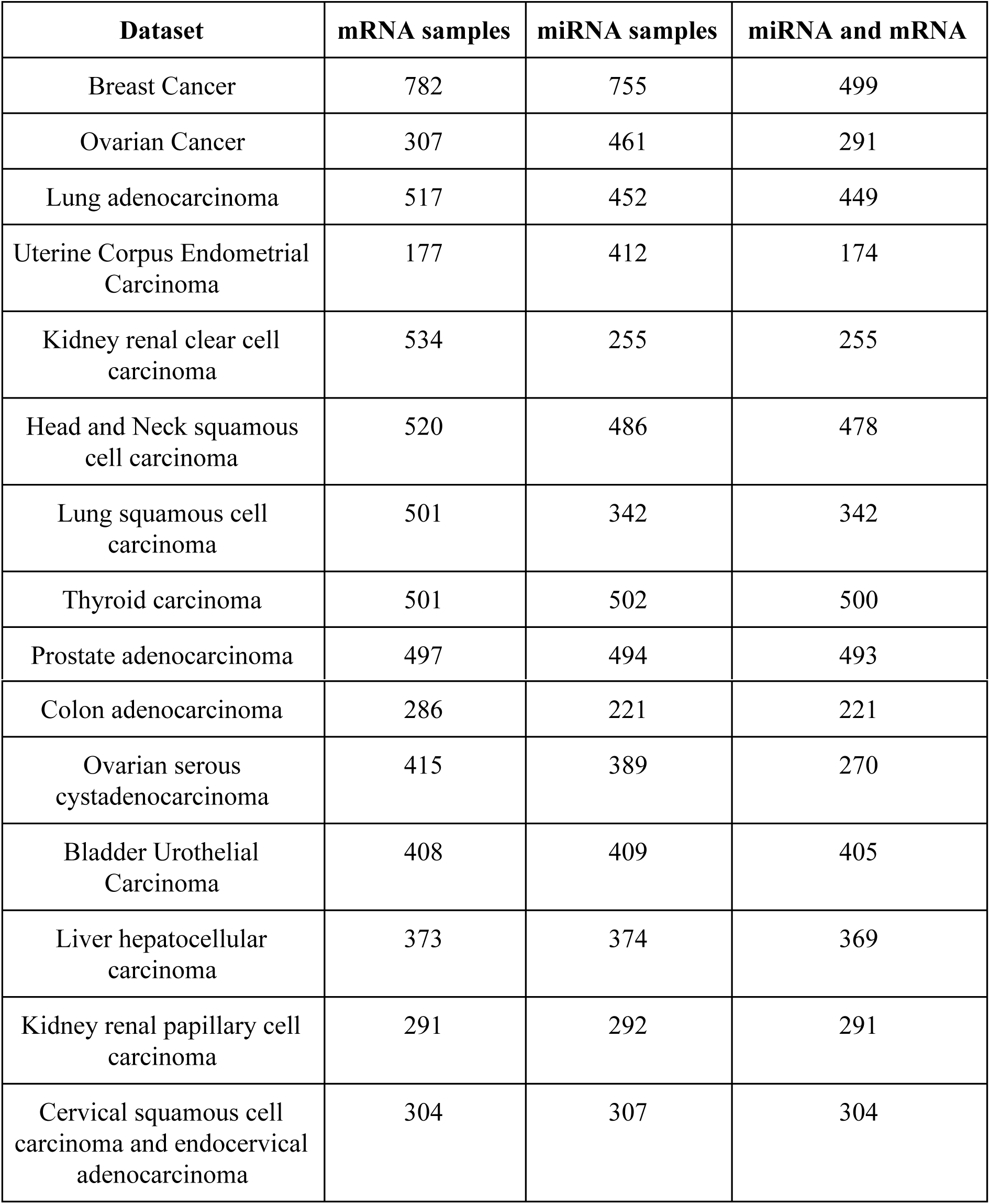
presents the breakdown of histologic subtypes in the TCGA dataset. All cancer types which were epithelial or glandular with respect to histology, and with at least 200 unique patient samples with paired mRNA and miRNA-sequencing data.

## References

1. Baumann M, Krause M, Overgaard J, Debus J, Bentzen SM, Daartz J, et al. Radiation oncology in the era of precision medicine. Nat Rev Cancer. 2016;16:234–49.

2. Scott JG, Berglund A, Schell MJ, Mihaylov I, Fulp WJ, Yue B, et al. A genome-based model for adjusting radiotherapy dose (GARD): a retrospective, cohort-based study. Lancet Oncol. 2017;18:202–11.

3. Curtis C, Shah SP, Chin S-F, Turashvili G, Rueda OM, Dunning MJ, et al. The genomic and transcriptomic architecture of 2,000 breast tumours reveals novel subgroups. Nature. 2012;486:346–52.

4. Cancer Genome Atlas Research Network, Weinstein JN, Collisson EA, Mills GB, Shaw KRM, Ozenberger BA, et al. The Cancer Genome Atlas Pan-Cancer analysis project. Nat Genet. 2013;45:1113–20.

5. Subramanian A, Tamayo P, Mootha VK, Mukherjee S, Ebert BL, Gillette MA, et al. Gene set enrichment analysis: a knowledge-based approach for interpreting genome-wide expression profiles. Proc Natl Acad Sci U S A. 2005;102:15545–50.

6. Barretina J, Caponigro G, Stransky N, Venkatesan K, Margolin AA, Kim S, et al. The Cancer Cell Line Encyclopedia enables predictive modelling of anticancer drug sensitivity. Nature. 2012;483:603–7.

7. Dhawan A, Scott JG, Harris AL, Buffa FM. Pan-cancer characterisation of microRNA with hallmarks of cancer reveals role of microRNA-mediated downregulation of tumour suppressor genes [Internet]. 2017. Available from: http://dx.doi.org/10.1101/238675

8. Piening BD, Wang P, Subramanian A, Paulovich AG. A radiation-derived gene expression signature predicts clinical outcome for breast cancer patients. Radiat Res. 2009;171:141–54.

9. Speers C, Zhao S, Liu M, Bartelink H, Pierce LJ, Feng FY. Development and Validation of a Novel Radiosensitivity Signature in Human Breast Cancer. Clin Cancer Res. 2015;21:3667–77.

10. Toustrup K, Sørensen BS, Nordsmark M, Busk M, Wiuf C, Alsner J, et al. Development of a hypoxia gene expression classifier with predictive impact for hypoxic modification of radiotherapy in head and neck cancer. Cancer Res. 2011;71:5923–31.

11. Eustace A, Mani N, Span PN, Irlam JJ, Taylor J, Betts GNJ, et al. A 26-Gene Hypoxia Signature Predicts Benefit from Hypoxia-Modifying Therapy in Laryngeal Cancer but Not Bladder Cancer. Clin Cancer Res. 2013;19:4879–88.

12. Lendahl U, Lee KL, Yang H, Poellinger L. Generating specificity and diversity in the transcriptional response to hypoxia. Nat Rev Genet. 2009;10:821–32.

13. Tawk B, Schwager C, Deffaa O, Dyckhoff G, Warta R, Linge A, et al. Comparative analysis of transcriptomics based hypoxia signatures in head- and neck squamous cell carcinoma. Radiother Oncol. 2016;118:350–8.

14. Manem VSK, Lambie M, Smirnov P, Kofia V, Freeman M, Koritzinsky M, et al. Modeling cellular response in large-scale radiogenomic databases to advance precision radiotherapy [Internet]. 2018. Available from: http://dx.doi.org/10.1101/449793

## REFERENCES

1. Haibe-Kains B, Desmedt C, Loi S, Culhane AC, Bontempi G, Quackenbush J, et al. A three-gene model to robustly identify breast cancer molecular subtypes. J Natl Cancer Inst. 2012;104:311–25.

2. Dhawan A, Barberis A, Cheng W-C, Domingo E, West C, Maughan T, et al. sigQC: A procedural approach for standardising the evaluation of gene signatures [Internet]. 2017. Available from: http://dx.doi.org/10.1101/203729

3. Piening BD, Wang P, Subramanian A, Paulovich AG. A radiation-derived gene expression signature predicts clinical outcome for breast cancer patients. Radiat Res. 2009;171:141–54.

4. Nuyten DSA, Kreike B, Hart AAM, Chi J-TA, Sneddon JB, Wessels LFA, et al. Predicting a local recurrence after breast-conserving therapy by gene expression profiling. Breast Cancer Res. 2006;8:R62.

5. Torres-Roca JF, Eschrich S, Zhao H, Bloom G, Sung J, McCarthy S, et al. Prediction of radiation sensitivity using a gene expression classifier. Cancer Res. 2005;65:7169–76.

6. Speers C, Zhao S, Liu M, Bartelink H, Pierce LJ, Feng FY. Development and Validation of a Novel Radiosensitivity Signature in Human Breast Cancer. Clin Cancer Res. 2015;21:3667–77.

7. Pitroda SP, Pashtan IM, Logan HL, Budke B, Darga TE, Weichselbaum RR, et al. DNA repair pathway gene expression score correlates with repair proficiency and tumor sensitivity to chemotherapy. Sci Transl Med. 2014;6:229ra42.

8. Kim HS, Kim SC, Kim SJ, Park CH, Jeung H-C, Kim YB, et al. Identification of a radiosensitivity signature using integrative metaanalysis of published microarray data for NCI-60 cancer cells. BMC Genomics. 2012;13:348.

9. de Jong MC, Ten Hoeve JJ, Grénman R, Wessels LF, Kerkhoven R, Te Riele H, et al. Pretreatment microRNA Expression Impacting on Epithelial-to-Mesenchymal Transition Predicts Intrinsic Radiosensitivity in Head and Neck Cancer Cell Lines and Patients. Clin Cancer Res. 2015;21:5630–8.

10. Amundson SA, Do KT, Vinikoor LC, Lee RA, Koch-Paiz CA, Ahn J, et al. Integrating global gene expression and radiation survival parameters across the 60 cell lines of the National Cancer Institute Anticancer Drug Screen. Cancer Res. 2008;68:415–24.

11. Abazeed ME, Adams DJ, Hurov KE, Tamayo P, Creighton CJ, Sonkin D, et al. Integrative radiogenomic profiling of squamous cell lung cancer. Cancer Res. 2013;73:6289–98.

12. Weichselbaum RR, Ishwaran H, Yoon T, Nuyten DSA, Baker SW, Khodarev N, et al. An interferon-related gene signature for DNA damage resistance is a predictive marker for chemotherapy and radiation for breast cancer. Proc Natl Acad Sci U S A. 2008;105:18490–5.

13. Tang Z, Zeng Q, Li Y, Zhang X, Suto MJ, Xu B, et al. Predicting radiotherapy response for patients with soft tissue sarcoma by developing a molecular signature. Oncol Rep. 2017;38:2814–24.

14. Toustrup K, Sørensen BS, Nordsmark M, Busk M, Wiuf C, Alsner J, et al. Development of a hypoxia gene expression classifier with predictive impact for hypoxic modification of radiotherapy in head and neck cancer. Cancer Res. 2011;71:5923–31.

15. Eustace A, Mani N, Span PN, Irlam JJ, Taylor J, Betts GNJ, et al. A 26-gene hypoxia signature predicts benefit from hypoxia-modifying therapy in laryngeal cancer but not bladder cancer. Clin Cancer Res. 2013;19:4879–88.

16. Lendahl U, Lee KL, Yang H, Poellinger L. Generating specificity and diversity in the transcriptional response to hypoxia. Nat Rev Genet. 2009;10:821–32.

17. Winter SC, Buffa FM, Silva P, Miller C, Valentine HR, Turley H, et al. Relation of a hypoxia metagene derived from head and neck cancer to prognosis of multiple cancers. Cancer Res. 2007;67:3441–9.

18. Buffa FM, Harris AL, West CM, Miller CJ. Large meta-analysis of multiple cancers reveals a common, compact and highly prognostic hypoxia metagene. Br J Cancer. 2010;102:428–35.

19. Hu Z, Fan C, Livasy C, He X, Oh DS, Ewend MG, et al. A compact VEGF signature associated with distant metastases and poor outcomes. BMC Med. 2009;7:9.

20. Sørensen BS, Toustrup K, Horsman MR, Overgaard J, Alsner J. Identifying pH independent hypoxia induced genes in human squamous cell carcinomas in vitro. Acta Oncol. 2010;49:895–905.

21. Yang L, Roberts D, Takhar M, Erho N, Bibby BAS, Thiruthaneeswaran N, et al. Development and Validation of a 28-gene Hypoxia-related Prognostic Signature for Localized Prostate Cancer. EBioMedicine. 2018;31:182–9.

22. Fjeldbo CS, Julin CH, Lando M, Forsberg MF, Aarnes E-K, Alsner J, et al. Integrative Analysis of DCE-MRI and Gene Expression Profiles in Construction of a Gene Classifier for Assessment of Hypoxia-Related Risk of Chemoradiotherapy Failure in Cervical Cancer. Clin Cancer Res. 2016;22:4067–76.

23. Yang L, Taylor J, Eustace A, Irlam JJ, Denley H, Hoskin PJ, et al. A Gene Signature for Selecting Benefit from Hypoxia Modification of Radiotherapy for High-Risk Bladder Cancer Patients. Clin Cancer Res. 2017;23:4761–8.

24. Ragnum HB, Vlatkovic L, Lie AK, Axcrona K, Julin CH, Frikstad KM, et al. The tumour hypoxia marker pimonidazole reflects a transcriptional programme associated with aggressive prostate cancer. Br J Cancer. 2015;112:382–90.

25. Seigneuric R, Starmans MHW, Fung G, Krishnapuram B, Nuyten DSA, van Erk A, et al. Impact of supervised gene signatures of early hypoxia on patient survival. Radiother Oncol. 2007;83:374–82.

26. Starmans MHW, Chu KC, Haider S, Nguyen F, Seigneuric R, Magagnin MG, et al. The prognostic value of temporal in vitro and in vivo derived hypoxia gene-expression signatures in breast cancer. Radiother Oncol. 2012;102:436–43.

27. Ghazoui Z, Buffa FM, Dunbier AK, Anderson H, Dexter T, Detre S, et al. Close and Stable Relationship between Proliferation and a Hypoxia Metagene in Aromatase Inhibitor-Treated ER-Positive Breast Cancer. Clin Cancer Res. 2011;17:3005–12.

28. van Malenstein H, Gevaert O, Libbrecht L, Daemen A, Allemeersch J, Nevens F, et al. A seven-gene set associated with chronic hypoxia of prognostic importance in hepatocellular carcinoma. Clin Cancer Res. 2010;16:4278–88.

